# metaMIC: reference-free Misassembly Identification and Correction of *de novo* metagenomic assemblies

**DOI:** 10.1101/2021.06.22.449514

**Authors:** Senying Lai, Shaojun Pan, Luis Pedro Coelho, Wei-Hua Chen, Xing-Ming Zhao

## Abstract

Evaluating the quality of metagenomic assemblies is important for constructing reliable metagenome-assembled genomes and downstream analyses. Here, we present metaMIC (https://github.com/ZhaoXM-Lab/metaMIC), a machine-learning based tool for identifying and correcting misassemblies in metagenomic assemblies. Benchmarking results on both simulated and real datasets demonstrate that metaMIC outperforms existing tools when identifying misassembled contigs. Furthermore, metaMIC is able to localize the misassembly breakpoints, and the correction of misassemblies by splitting at misassembly breakpoints can improve downstream scaffolding and binning results.

## Background

Constructing reliable metagenome-assembled genomes (MAGs) is of great importance for understanding microbial communities and downstream functional analysis, such as taxonomic annotations and reconstruction of metabolic processes [1–4]. MAGs are obtained by binning assembled contigs into bins, the quality of which can be significantly affected by the assembly errors in contigs. For example, the chimerical assemblies consisting of two or more genomes can introduce contamination for reconstructed MAGs, potentially resulting in misleading biological conclusions [5]. Despite the progress in assembly algorithms, errors are still prevalent in metagenomic-assembled contigs owing to the inherent complexity of metagenomic data. Assembly errors including inter- and intra-species misassemblies are caused by repetitive genomic regions that occur within the same genome or conserved sequences shared among distinct organisms, which is especially likely to happen when multiple closely-related strains are present in the same environment [6, 7]. Therefore, the evaluation of metagenomic assemblies is critical for constructing high-quality and reliable MAGs.

Approaches that have been proposed for assessing the quality of metagenomic assemblies can be grouped into two categories: *reference-based* and *reference-free* approaches. Reference-based methods evaluate the *de novo* assemblies by aligning them against corresponding reference genomes. For example, MetaQUAST [8], the metagenomic-adapted version of QUAST [9], detects misassemblies such as translocation, inversion and relocation by mapping the metagenomic contigs to a set of closely-related reference genomes. However, it is difficult to distinguish errors from real structural variation. Moreover, reference genomes are available for only a small fraction of organisms found in real environments, which limits these approaches to previously-sequenced species [10]. Therefore, the evaluation of metagenomic assemblies would benefit from reference-free methods. Typically these methods exploit features such as the high variation in coverage depth or inconsistent insert distance of paired-end reads to indicate possible repeat collapse, misjoins or error insertions/deletions [11]. Popular reference-free methods include ALE [12], DeepMAsED [13], SuRankCo [14] and VALET [15]. ALE measures the quality of assemblies as the likelihood that the observed reads are generated from a given assembly by modeling the sequencing process. SuRankCo uses a machine learning approach to provide quality scores for contigs based on characteristics of contigs such as length and coverage. VALET detects misassemblies based on the combination of multiple metrics extracted from the alignment of reads to contigs. DeepMAsED employs a deep learning approach to identify misassembled contigs. Despite the great value of those approaches for evaluating metagenomic assembly quality, only VALET and ALE predict the position where the misassembly errors are introduced and none of these methods have functionality for correcting metagenomic misassemblies. More importantly, VALET and SuRankCo are no longer maintained, and software incompatibilities hinder their use.

Here, we present a novel tool called metaMIC which performs reference-free misassembly identification and correction in *de novo* metagenomic assemblies. metaMIC can identify misassembled contigs, localize misassembly breakpoints within misassembled contigs and then correct misassemblies by splitting misassembled contigs at breakpoints. Benchmarking results on both simulated and real metagenomics data show that metaMIC can identify misassembled contigs with higher accuracy than state-of-the-art tools, and precisely localize the misassembly error regions and recognize breakpoints in both single genomic and metagenomic assemblies. By comparing the scaffolding and binning results before and after metaMIC correction, we demonstrate that the correction of misassemblies by metaMIC can improve the scaffolding and binning results.

## Results

### Overview of the metaMIC pipeline

metaMIC is a fully automated tool for identifying and correcting misassemblies in metagenomic contigs using the following three steps (Fig. 1). First, various types of features were extracted from the alignment between paired-end sequencing reads and each contig, including sequencing coverage, nucleotide variants, mate pair consistency, and *k*-mer abundance differences (KAD) [16] between mapped reads and the contig. The KAD method was previously developed for evaluating the accuracy of nucleotide base quality in single genomic assemblies. Here, we extended KAD to metagenomic assemblies to measure the overall consistency between mapped reads and corresponding contigs (see Methods). Secondly, the features extracted in the first step are used as input to a random forest classifier for identifying misassembled contigs, where the classifier is trained with simulated bacterial metagenomic communities to discriminate misassembled contigs from correctly assembled ones. Thirdly, metaMIC will localize misassembly breakpoint(s) in each misassembled contig, namely the point at which the left and right flanking sequences are predicted to have originated from different genomes or locations. As most misassemblies are chimeras where two fragments from different locations or with different orientations are mistakenly connected and not just random sequences being generated [9], misassemblies can be corrected by breaking up the contigs into two (or more) correctly assembled contigs.

**Fig. 1.**
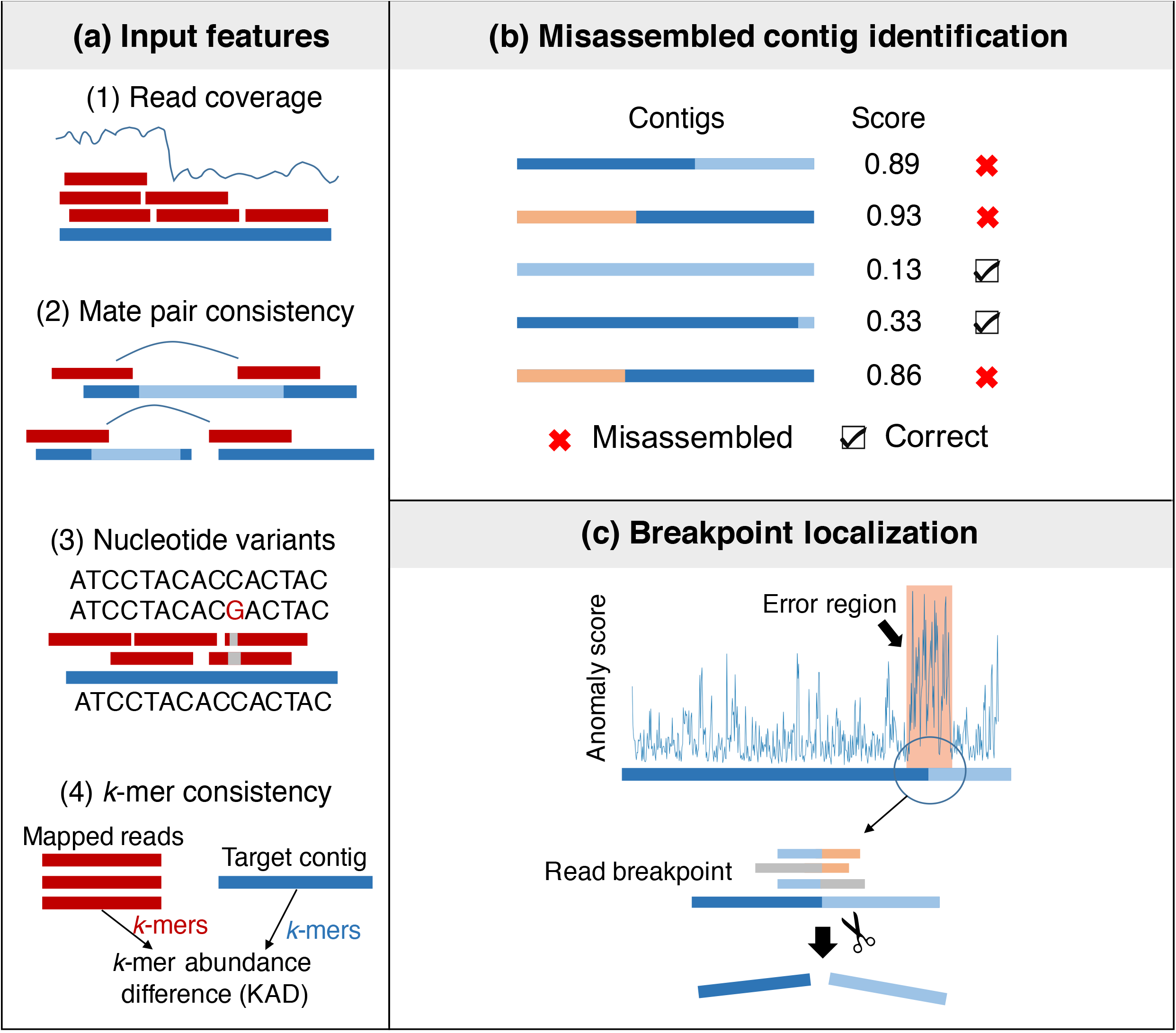
Overall framework of metaMIC. **a** metaMIC extracts four types of features from the alignment of paired end reads to contigs: read coverage, nucleotide variants, mate pair consistency, and *k*-mer abundance consistency. **b** Misassembled contigs are identified by metaMIC based on the four features. **c** metaMIC first identifies the error regions containing misassembly breakpoints, and then recognizes the exact positions of breakpoints and corrects misassemblies by splitting misassembled contigs at breakpoints.

### Identifying misassembled contigs in simulated metagenomic datasets

To evaluate metaMIC, we tested it on simulated metagenomic datasets obtained from CAMI (the Critical Assessment of Metagenome Interpretation) [2] that comprise a known mixture of organisms. We first evaluated metaMIC on the Medium (*CAMI1-Medium*) and High-diversity communities (*CAMI1-High*) to see how dataset complexity will influence the accuracy of metaMIC. We noticed that the types of misassemblies identified in these two datasets were slightly different, and the CAMI1-High dataset contains more inter-species translocations and higher proportion of misassemblies while the CAMI1-Medium dataset contains more relocations (see Figs. S1, S2), which is consistent with previous conclusion that datasets with higher intra-species diversity tend to have more inter-species translocation misassemblies [13]. Compared with CAMI1-High metaMIC performed better on (Fig. 2a; although still significantly better than existing tools) CAMI1-Medium, implying that the higher microbial diversity increases the challenge of identifying misassembled contigs. We further compared metaMIC on these datasets against ALE [12] and DeepMAsED [13] (See Methods). As shown in Fig. 2a, metaMIC significantly outperforms both ALE and DeepMAsED on the two datasets, as MetaMIC achieved 4-fold higher AUPRC (area under the precision-recall curve).

**Fig. 2.**
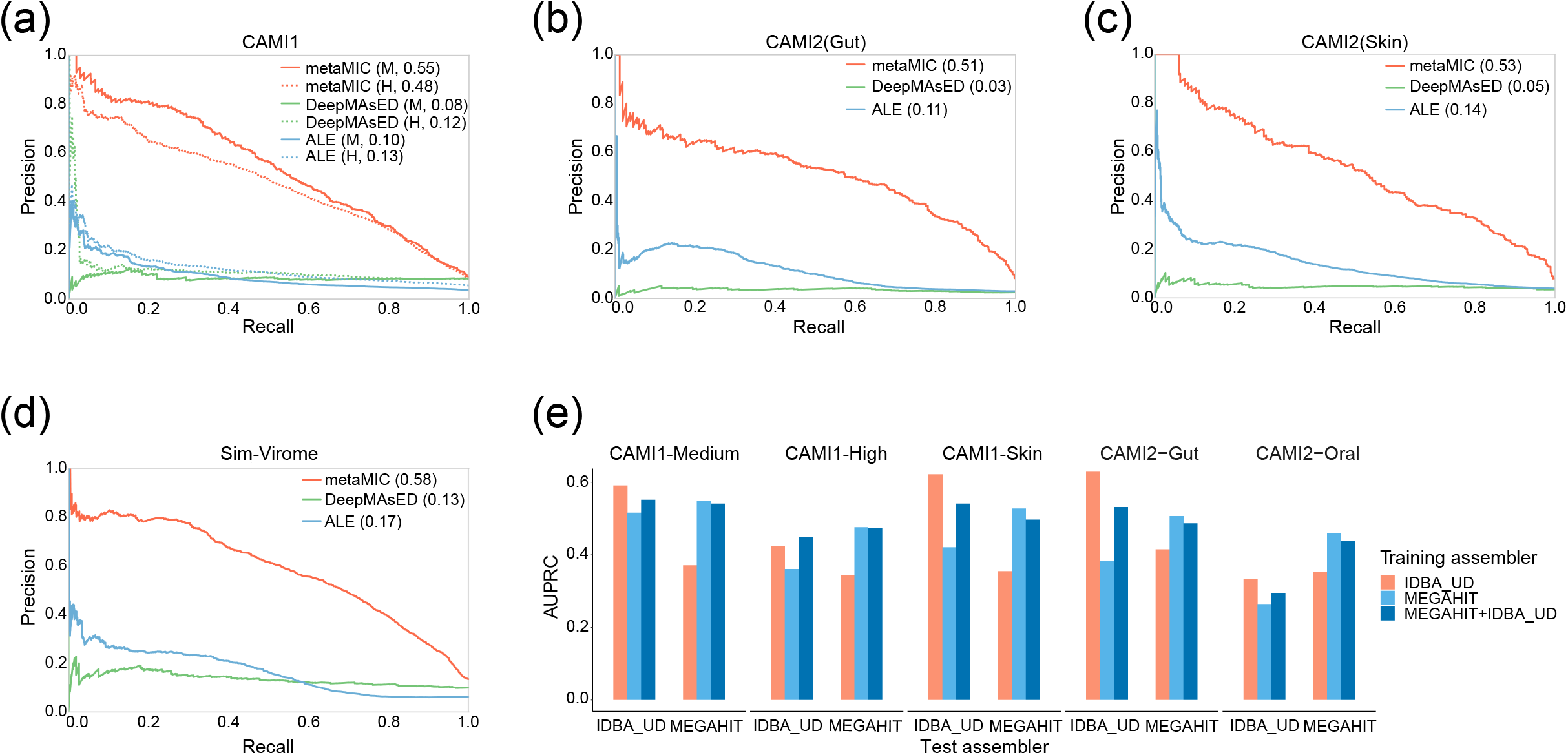
metaMIC outperforms ALE and DeepMAsED in identifying misassembled contigs in simulated metagenomic datasets. **a-d** The performance of the three tools on the CAMI-medium (M) and high-complexity (H) communities (**a)**, *CAMI2-Skin* (**b**), *CAMI2-Gut* (**c**), and simulated virome dataset (*Sim-Virome*) (**d**). **e** The AUPRC scores of metaMIC on test datasets assembled by MEGAHIT or IDBA_UD (Test assembler), where metaMIC were trained on contigs from training datasets assembled by MEGAHIT, IDBA_UD, or jointly by MEGAHIT and IDBA_UD (MEGAHIT+IDBA_UD).

We also evaluated metaMIC and other tools on simulated metagenomic datasets from three different human body sites: gastrointestinal tract (*CAMI2-Gut*), skin (*CAMI2-Skin*) and oral cavity (*CAMI2-Oral*). As shown in Figs. 2b, c and Fig. S3, metaMIC has the highest precision when identifying misassembled contigs at any recall threshold. Additionally, we tested metaMIC on a simulated virome datasets (*Sim-Virome*), which were simulated based on 1,000 complete viral genomes randomly selected from NCBI RefSeq collection [17] (See Methods). The Sim-Virome contains mainly translocations and relocations but few inter-species translocations and inversions. We found that metaMIC still significantly outperforms both ALE and DeepMAsED on Sim-Virome dataset as shown in Fig. 2d, indicating that metaMIC can also be used for virome assemblies besides bacterial metagenomic assemblies.

As metaMIC can be trained on contigs assembled by different assemblers, we further investigated the impact of different assemblers on the performance of metaMIC when identifying misassembled contigs. Here, two popular assemblers, i.e. MEGAHIT and IDBA_UD, used for metagenomic data were considered. As shown in Fig. 2e, we found that metaMIC performed best when it was trained on the same assembler as it was later evaluated. Therefore, we recommend to use metaMIC trained on the contigs generated by the same assembler. For version 1.0, metaMIC provides builtin models supporting MEGAHIT and IDBA_UD as well as the ability to generate new models based on the assembler specified by users.

### metaMIC can identify breakpoints with higher accuracy in misassembled contigs

Beyond identifying misassembled contigs, metaMIC is able to accurately recognize the misassembly breakpoints, at which the misassembled contigs can be split into shorter ones. From the distribution, we can clearly see that the error regions containing any misassembly type generally have significantly higher anomaly scores than error-free regions, and the inter-species translocation error is most prevalent in the dataset. In the CAMI datasets, it is indeed the inter-species translocation error that occurs most often (Fig. S2). The differential distribution of anomaly scores between error and error-free regions implies that the anomaly score has the potential to recognize the error regions. We also noticed that the misassembly sites are usually read breakpoints (locations at which the boundaries of aligned read fragments do not coincide with the ends of corresponding reads) [18]. Similar to anomaly scores, we found that the read breakpoint ratio was significantly different between error regions and error-free regions (Fig. 3b, see also Figs. S7, S8).

**Fig. 3.**
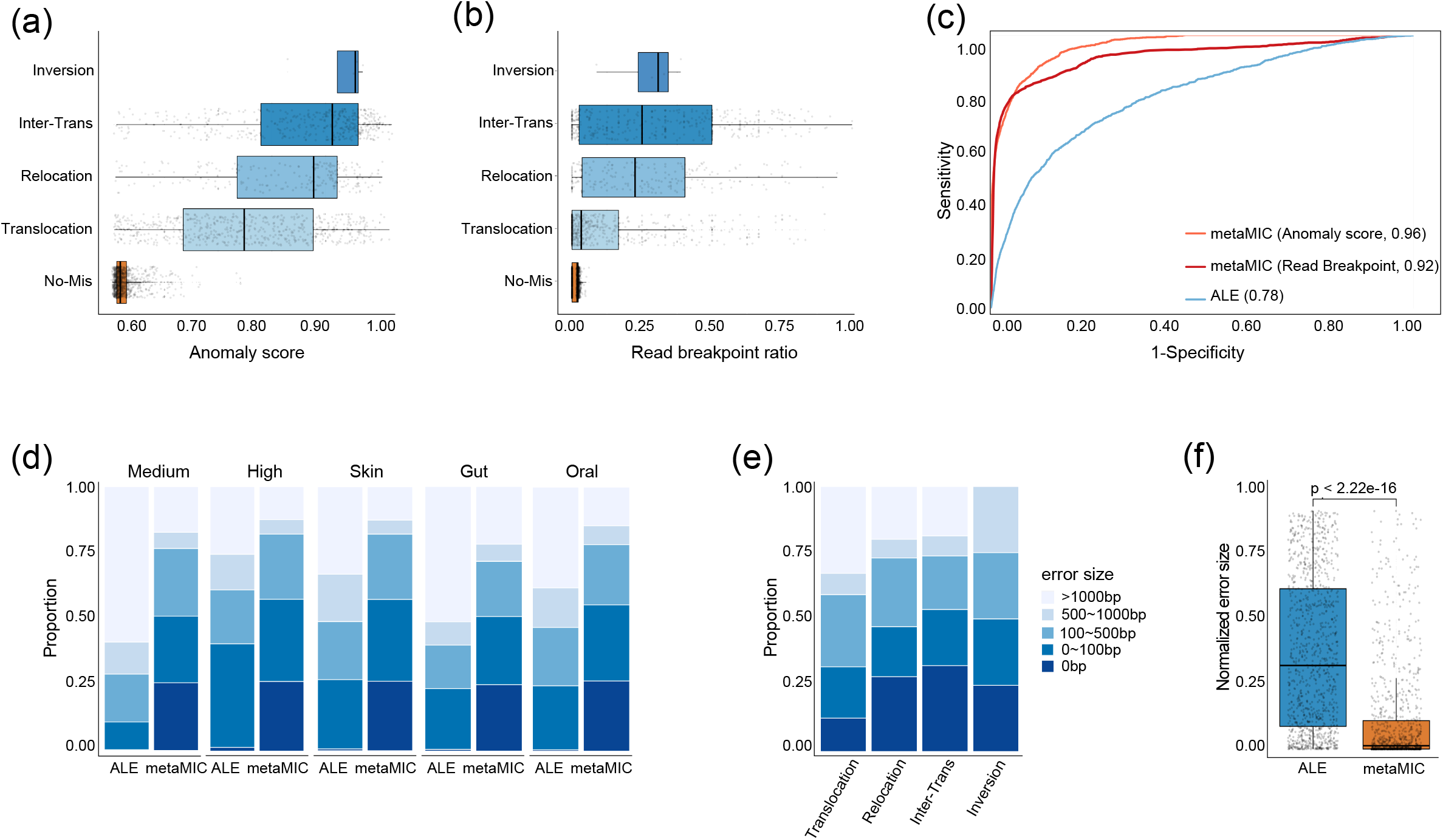
The performance of metaMIC in localizing misassembly breakpoints on CAMI datasets. **a, b** The distribution of anomaly scores (**a**) and read breakpoint ratios (**b**) of different misassembly types across contigs from CAMI1-Medium. **c** The receiver operation curves by ALE, anomaly scores and read breakpoint ratios when discriminating error regions from error-free regions in CAMI1-Medium, respectively. **d, e** The distribution of error size of misassembly breakpoints recognized by metaMIC on CAMI1-Medium (*Medium*), CAMI1-High (*High*), CAMI2-Skin (*Skin*), CAMI2-Gut (*Gut*) and CAMI2-Oral (*Oral*) (**d**), and different misassembly types in CAMI1-Medium (**e**). **f** The distribution of normalized error size of misassembly breakpoints recognized by metaMIC and ALE on CAMI1-Medium.

Due to the potential of read breakpoint ratio and anomaly score to localize the error regions, we want to see whether metaMIC can use these two features to separate the erroneous regions from error-free regions. From the receiver operation curves shown in Fig. 3c, we can see that with either anomaly score or read breakpoint ratio, metaMIC can classify the error regions containing misassembly breakpoints with error-free regions more accurately than ALE. To combine the usages of these two features, metaMIC first localizes the error regions in a misassembled contigs with the help of anomaly score, and then identifies the exact breakpoints in an error region based on the read breakpoint ratio.

We evaluated the performance of both metaMIC and ALE on the five datasets from CAMI as shown in Fig. 3d. We observed that approximately 71-86% of the metaMIC-predicted breakpoints were within 500bp compared to 26-48% of those by ALE. More importantly, metaMIC could predict the exact locations for ~25% of the breakpoints with the use of read breakpoints. Again, inter-species translocations or inversions can be detected with higher accuracy relative to other misassembly types (Fig. 3e), consistent with previous results that they were supported by more fragmentally aligned reads and had higher anomaly scores as compared with other error types (see Figs. 3a, b; Fig. S8). Given the possible influence of contig length on the prediction error size, we normalized the error size by the contig length, and compared the results of metaMIC with those of ALE. As shown in Fig. 3f, metaMIC still significantly outperforms ALE with respect to the normalized error size (Wilcoxon test, p-value <2.22e-16), where the median and mean of the metaMIC’s normalized error size were 0.01 and 0.11, respectively, compared to 0.39 and 0.34 for ALE (see also Fig. S9).

### Splitting misassembled contigs improves downstream binning performance

As metaMIC can identify breakpoints in misassembled contigs, it can split misassembled contigs at breakpoints and reduce the number of misassemblies (see Methods); although the contiguity could be slightly decreased due to more fragmented contigs [19]. To see how the correction of splitting misassembled contigs at breakpoints employed by metaMIC will influence downstream analyses, we binned the contigs in the simulated datasets. We then assessed the binning performance over the original and metaMIC-corrected contigs by counting the number of obtained high-quality bins. We can see in Fig. 4a that metaMIC correction increases the number of near-complete reconstructed bins (completeness above 90%, contamination below 5% [3]) by 10-20% (see also Fig. S13a, Table S1), showing that the correction of metagenomic miassemblies has significant impact on downstream binning. We noticed that most of the misassemblies corrected by metaMIC were inter-species translocations that were also the main sources of chimeras and assembly errors in CAMI datasets (Fig. S2, Table S2). From Fig. 4b and also those shown in Fig. S13b, we can see that bin-wise F1 scores of those bins constructed from corrected contigs are significantly improved compared with the results over original contigs, indicating that the reconstructed bins can better represent the reference genomes after metaMIC correction. The above results clearly demonstrate that the correction of metagenomic misassemblies by metaMIC can significantly improve the resulting bins in term of both completeness and contaminations, which is important for understanding the complex microbiota communities.

**Fig. 4.**
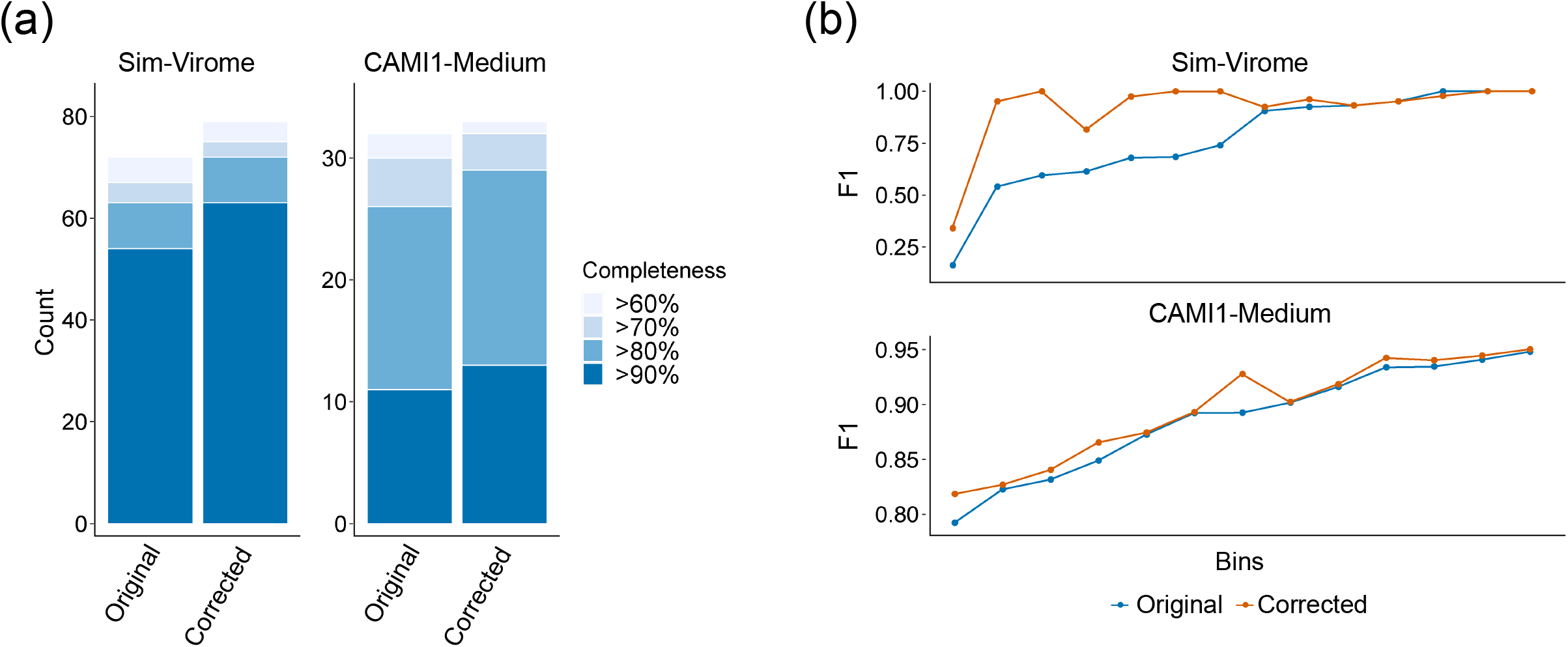
Splitting misassembled contigs at breakpoints improves the downstream binning results over *Sim-Virome* and *CAMI1-Medium* datasets. **a** The number of high-quality bins with low contamination (<5%) of different completeness reconstructed from original and corrected contigs. **b** The distribution of F1 scores for bins reconstructed based on contigs before and after correction, where only those bins whose results change before and after correction were shown for clearness.

### Application of metaMIC to real metagenomic datasets

To better evaluate the performance of metaMIC, we applied metaMIC to two recent human gut metagenomics datasets from Ethiopian [20] and Madagascar [21] cohorts that consist of 50 and 112 samples, respectively. In total, metaMIC respectively identified 5,905 and 18,436 misassembled contigs in *Ethiopian* and *Madagascar* datasets, which represents 2.59% and 4.53% of all contigs in the two datasets. We then separately binned the original and corrected contigs into bins. Strikingly, we found that ~20% of the resulting original-bins contained misassemblies, although the latter accounted for less than 5% of all contigs (See Table S3). As previous results have demonstrated that metaMIC correction can improve the binning results in simulated datasets, we further explored whether the correction step employed by metaMIC can improve downstream results in real datasets. As shown in Fig. 5, in addition to obtaining more bins of high quality (Completeness >90 and Contamination <5) (Fig. 5a, Table S3), the corrected bins had an equal or higher F1 score compared to the corresponding original bins (Fig. 5b). The results indicate that the misassembled contigs identified by metaMIC in these two real datasets are really misassembled, the correction of which can significantly improve downstream analysis results.

**Fig. 5.**
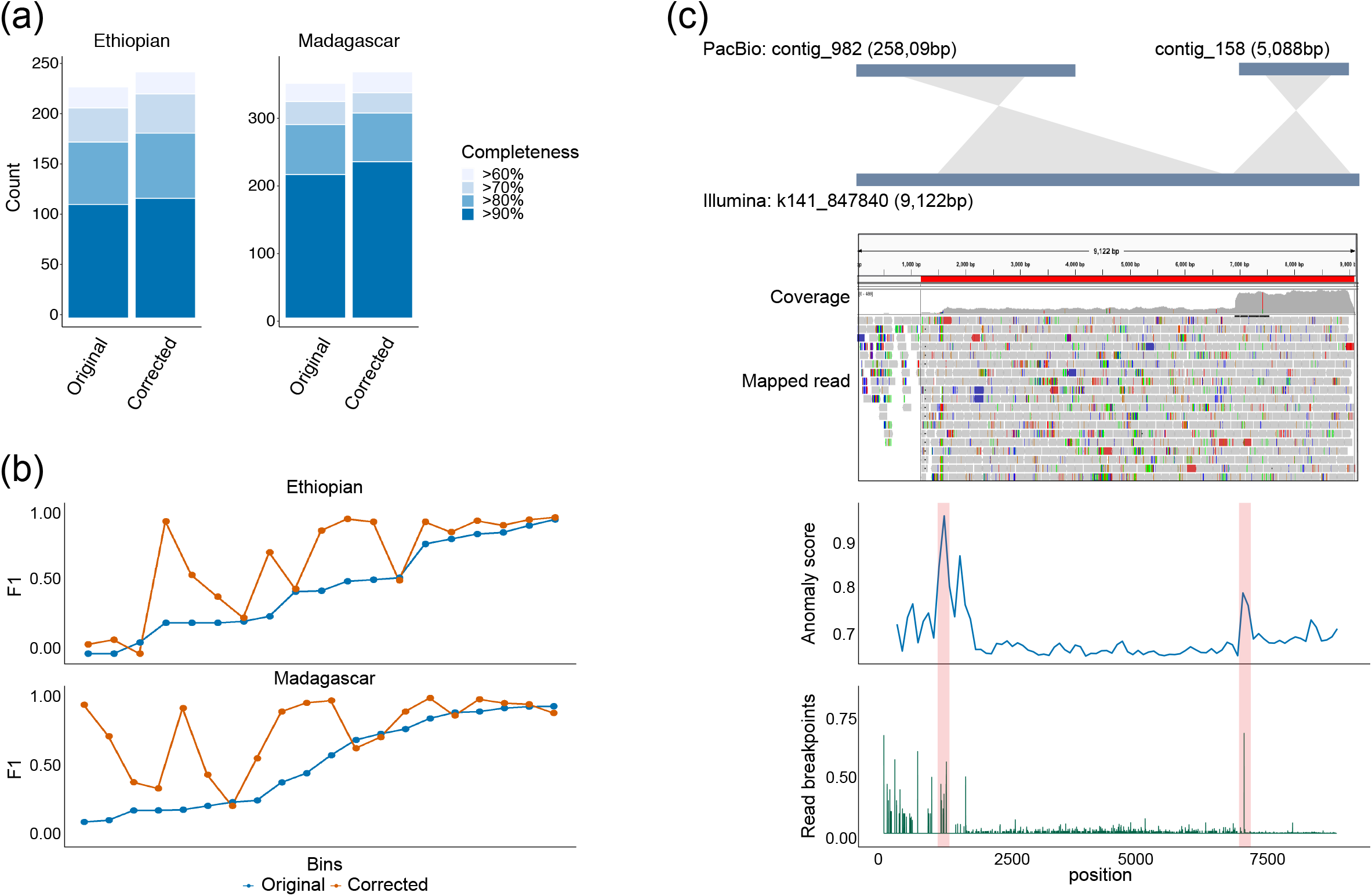
The performance of metaMIC on real metagenomic datasets. **a** The number of bins of different completeness with low contamination (<5%) reconstructed from original and corrected assemblies of ‘*Ethiopian*’ (left) and ‘*Madagascar*’ (right) cohorts. **b** Comparison of F1 scores for reconstructed bins before and after correction of contigs from ‘*Ethiopian*’ (top) and ‘*Madagascar*’ (bottom) cohorts. **c** An example of a predicted misassembled contig “k141_847840” assembled from combined rumen fluid and solid sample. The top plot shows the alignment result of Illumina short-read assembled contig “k141_847840” and PacBio long-read assembled contigs (“contig_982” and “contig_158”), where two regions in the “k141_84780” (1201-6738bp and 6920-8700bp) were aligned to “contig_982” and “contig_158”, respectively. The middle figure shows a snapshot of Integrative Genomics Viewer for contig “k141_847840”. The bottom plot shows the anomaly score (blue) and read breakpoint ratio (green) across contig “k141_847840”.

As these contamination metrics are based on *in silico* evaluation, we further tested the ability of identifying misassemblies using another metagenomic dataset (a combined rumen fluid and solid sample) where both short and long reads are available. Since the long reads from PacBio platforms are able to span repeats [22, 23], which are the main contributor to misassemblies, we can therefore use the long-read assemblies as gold standards to validate our predicted misassembled contigs from the short-read assemblies. In total, metaMIC identified 692 misassembled contigs (approximately 2.5%) in the short-read assemblies. By manual inspection of the alignments between PacBio assemblies and short-read assemblies, we can validate a subset of metaMIC predictions (Fig. 5c and Fig. S14). For instance, there exist two peaks at positions of 1200bp and 6920bp in the contig of “k141_847840” according to the anomaly scores by metaMIC, and both peaks, especially the one at 6920bp, contain higher read breakpoint counts implying possible misassembly breakpoints at these two locations. When aligning this contig against the long-read assemblies, we found that two regions in this contig (1201-6738bp and 6920-8700bp) were indeed aligned to two different long-read assembled contigs, and a change-point in the read coverage at 6920bp can be observed (see Fig. 5c), indicating that there are actually two contigs wrongly assembled into one contig at position of 6920bp. We also found that only a few reads can be aligned to the region of 0-1200bp, suggesting this region may be extended mistakenly by the assembler.

### Application of metaMIC to isolate genomes

Since metaMIC can identify and correct intra-species misassemblies such as inversions and relocations, metaMIC can also be applied to isolate genomes. We tested metaMIC on four real datasets from GAGE-B project [24], which aimed to evaluate assembly algorithms on isolate genomes. We tested metaMIC on *B. cereus, M. abscessus, R. sphaeroides* and *V. cholerae*, where the raw reads, assembled contigs [25] and curated reference genomes are available for these four species. metaMIC was ran on the assemblies downloaded from GAGE-B project and its performance was evaluated with the results by QUAST [9] as gold standard. These four datasets contain mainly relocations but a few translocations (Table S4). We noticed that similar to metagenomes, the error regions in isolate genomes also have higher anomaly scores and more read breakpoints than error-free regions (Fig. S15). We then compared metaMIC against MEC [26], a recently-developed misassembly correction tool, when identifying misassembly breakpoints on the four isolate genomes. As shown in Table 2, metaMIC identified more true misassemblies than MEC, where approximately 80% misassemblies can be corrected compared to ~30% of MEC; and after the correction by metaMIC, the total number of bases of uncorrected misassembled contigs (i.e. misassembled contig length in Table 2) was significantly reduced compared with that by MEC.

**Table 1.** Performance comparison of metaMIC and MEC on four real datasets from the GAGE-B project.

| Species | Correction tool | Misassembled<br>contig length | FN | TP | FP |
| --- | --- | --- | --- | --- | --- |
| M. abscessus | raw | 1,189,973 | 20 | \ | \ |
|  | MEC | 982,986 | 14 | 6 | 2 |
|  | metaMIC | 593,741 | 6 | 14 | 2 |
| V. cholerae | raw | 597,777 | 7 | \ | \ |
|  | MEC | 597,183 | 6 | 1 | 0 |
|  | metaMIC | 205,644 | 3 | 4 | 1 |
| R. sphaeroides | raw | 135,153 | 2 | \ | \ |
|  | MEC | 64,489 | 1 | 1 | 0 |
|  | metaMIC | 0 | 0 | 2 | 7 |
| B. cereus | raw | 117,830 | 5 | \ | \ |
|  | MEC | 56,086 | 4 | 1 | 0 |
|  | metaMIC | 28,068 | 1 | 4 | 3 |

Misassembled contig length denotes the total number of bases in the raw misassembled contigs or the misassembled contigs that cannot be corrected by MEC or metaMIC; True positive (TP) is the number of true misassemblies identified by the error correction tool; False positive (FP) is the number of misassemblies which are actually correct but mistakenly identified as misassemblies; False negative (FN) denotes the number of true misassemblies that are not identified.

**Table 2.**
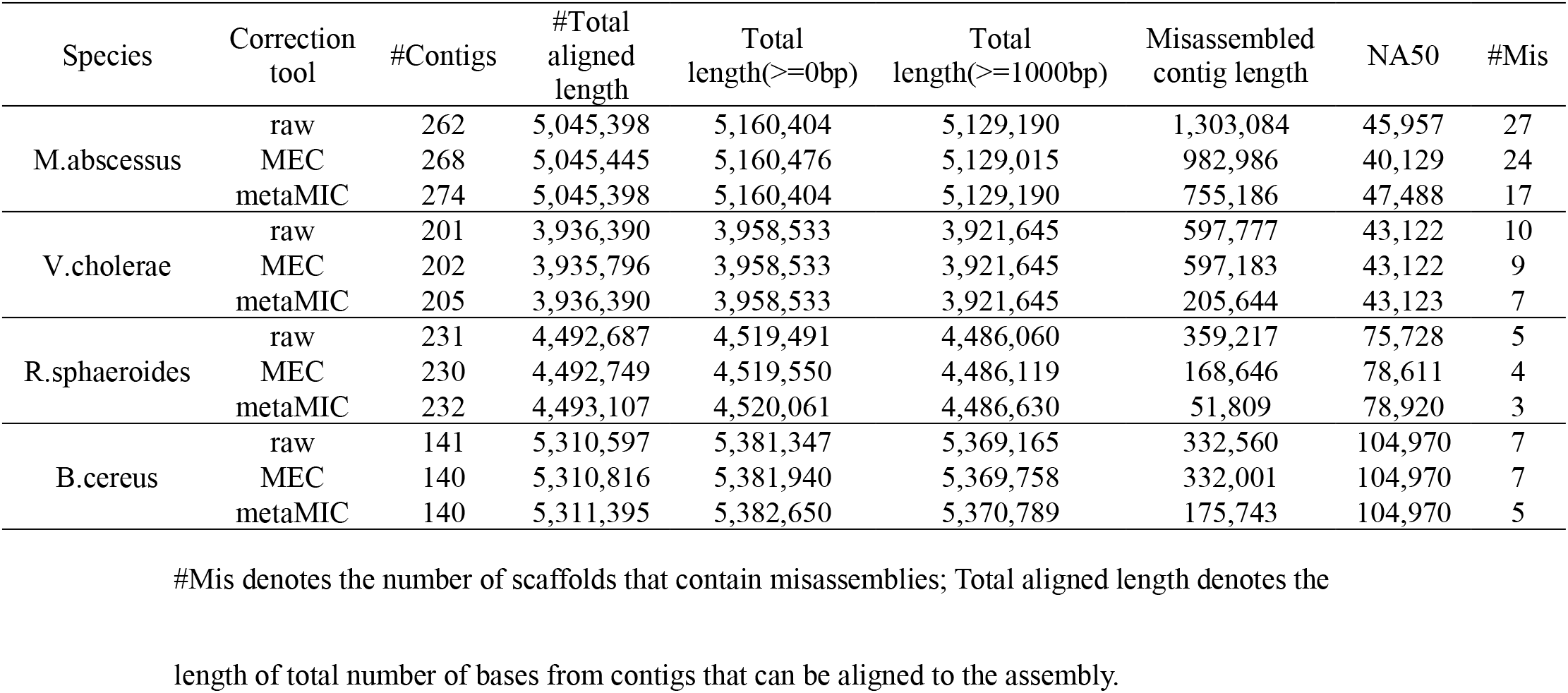
Comparison of BESST scaffolding results of contigs before and after correction.

| Species | Correction tool | #Contigs | #Total aligned length | Total length(>=0bp) | Total length(>=1000bp) | Misassembled contig length | NA50 | #Mis |
| --- | --- | --- | --- | --- | --- | --- | --- | --- |
| M.abscessus | raw | 262 | 5,045,398 | 5,160,404 | 5,129,190 | 1,303,084 | 45,957 | 27 |
|  | MEC | 268 | 5,045,445 | 5,160,476 | 5,129,015 | 982,986 | 40,129 | 24 |
|  | metaMIC | 274 | 5,045,398 | 5,160,404 | 5,129,190 | 755,186 | 47,488 | 17 |
| V.cholerae | raw | 201 | 3,936,390 | 3,958,533 | 3,921,645 | 597,777 | 43,122 | 10 |
|  | MEC | 202 | 3,935,796 | 3,958,533 | 3,921,645 | 597,183 | 43,122 | 9 |
|  | metaMIC | 205 | 3,936,390 | 3,958,533 | 3,921,645 | 205,644 | 43,123 | 7 |
| R.sphaeroides | raw | 231 | 4,492,687 | 4,519,491 | 4,486,060 | 359,217 | 75,728 | 5 |
|  | MEC | 230 | 4,492,749 | 4,519,550 | 4,486,119 | 168,646 | 78,611 | 4 |
|  | metaMIC | 232 | 4,493,107 | 4,520,061 | 4,486,630 | 51,809 | 78,920 | 3 |
| B.cereus | raw | 141 | 5,310,597 | 5,381,347 | 5,369,165 | 332,560 | 104,970 | 7 |
|  | MEC | 140 | 5,310,816 | 5,381,940 | 5,369,758 | 332,001 | 104,970 | 7 |
|  | metaMIC | 140 | 5,311,395 | 5,382,650 | 5,370,789 | 175,743 | 104,970 | 5 |

To further see influence of misassembly correction on isolate genomes, we scaffolded original and corrected contigs separately with popular scaffolders including BESST [27] and ScaffMatch [28], and then used QUAST to evaluate the scaffolding results. As seen in Table 3 and supplementary Table 6, the number of misassemblies in the scaffolding results based on metaMIC’s corrected contigs was much lower than that based on the original uncorrected contigs, and metaMIC significantly outperforms MEC in terms of misassembled contig length. Moreover, metaMIC performs comparably well or better compared against MEC in terms of NA50 and total aligned length, and performs better especially for *R.sphaeroides*. The above results clearly show the effectiveness of metaMIC when identifying and correcting misassembled contigs on isolate genomes, and also the capability of maintaining or improving the contiguity of downstream scaffolding after correction.

## Discussion

We present a novel tool named metaMIC to identify and correct misassembled contigs from *de novo* metagenomic assemblies and demonstrate its effectiveness on both simulated and real metagenomic datasets of varying complexity. Unlike most existing metagenomic assembly evaluation methods that only evaluate individual contigs or overall assemblies, metaMIC is capable of localizing the misassembly breakpoints and then corrects the misassembled contigs at breakpoints. By integrating various types of features extracted from both reads and assemblies, including read coverage, mate pair consistency, nucleotide variants and *k*-mer abundance consistency, metaMIC is able to detect intra- and inter-species misassemblies. Additionally, metaMIC can also be applied on isolate genomes given its ability in identifying intra-species misassemblies. After the correction of misassemblies, metaMIC can significantly help improve the performance of downstream analysis including binning and scaffolding.

In this study, the performance of metaMIC is mainly shown on the metagenomic assemblies assembled by MEGAHIT due to its high memory efficiency [29]. As different assemblers tend to be biased to certain types of misassemblies, the models trained on the outputs of one assembler may not transfer well to other assemblers. Note that metaMIC can be easily extended to work on the metagenomic assemblies by other assembler tools if the training datasets generated by the corresponding assemblers are available. We suggest to use metaMIC on the datasets from the same assembler as the one it is trained on.

metaMIC scans each contig with a sliding window of 100bp to localize the candidate error regions. Generally, a shorter window size can have a higher resolution to pinpoint error regions but require more computation resources while the longer window size can be robust to noise but are more likely to cover multiple errors. In addition, metaMIC currently cannot distinguish the types of assembly errors. In the future, more work is needed to determine the error types which in turn can help to correct misassemblies more accurately.

metaMIC correction mainly relies on splitting contigs at misassembly breakpoints. However, caution should be needed here as more fragmented sequences will be generated and mistakenly splitting may result in disrupted gene structure, which can have adverse influence on downstream functional genomic analysis. Although we have showed that metaMIC correction can improve the downstream binning results, the quality of reconstructed draft bins can be further improved if the broken contigs can be joined into scaffolds correctly. Thus, the combination of metaMIC and scaffolding algorithms will be a promising direction for future research, leading to effective approaches for reconstructing genomes from sequencing data with higher quality and completeness.

Several directions hold promise for further improvements to metaMIC. Firstly, metagenomic read mapping can be evaluated in more robust manner by aligning multi-assigned reads in a probabilistic manner to their contig of origin [30] or using base-level quality metrics such as CIGAR strings [31]. Secondly, increasing the amount of training data and integrating other assemblers such as metaSPAdes [32] are also potential directions for the improvement of metaMIC. Thirdly, the factors that may result in false positive predictions, such as structural variation within species of high similarity and G-C bias in sequencing coverage could be taken into account in future work. Finally, as reference genomes of many bacterium are available, a better performance can be achieved by the combination of reference-free and reference-based approaches.

## Conclusions

Here, a novel tool named metaMIC is developed for identifying and correcting misassemblies in *de novo* metagenomic assemblies without the use of reference genomes. Benchmarking on both simulated and real datasets, we show that metaMIC is able to pinpoint misassemblies in both single and metagenomic assemblies. We also demonstrate that metaMIC is able to improve the scaffolding or binning results by splitting misassembled contigs at misassembly breakpoints. As none of current assemblers can achieve a completely accurate assembly and misassemblies in contigs have negative influence on downstream analysis, we expect metaMIC can serve as a guide in optimizing metagenomic assemblies and help researchers be aware of problematic regions in assembled contigs, so as to avoid misleading downstream biological analysis.

## Methods

### metaMIC workflow

metaMIC is implemented in Python3 (Python ≥ 3.6). It requires assembled contigs in FASTA format and paired-end reads in FASTA or FASTQ format as input. Alternatively, the user can provide a BAM file with read pairs mapping to contigs. Given the contigs, metaMIC will first identify the misassemblies by employing a random forest classifier trained on the features extracted from reads and contigs. Next, metaMIC will identify the regions containing misassembly breakpoints in the misassembled contigs based on the anomaly scores, and then recognize the exact positions of the breakpoints in the error regions. Then metaMIC will correct the misassemblies by splitting the contigs at the breakpoints. The details will be given below.

#### Features extracted from reads and contigs

BWA-MEM (v.0.7.17) [33] is used to map paired-end reads to assemblies, followed by using samtools (v1.9) [34] to filter low quality mappings and sorting the alignments. Then four types of features will be extracted from the sorted BAM file, including read coverage, mate-pair consistency, nucleotide variants and *k*-mer abundance difference.

For each paired-end reads with left and right mate reads, the insert size corresponding to the distance between two mates is assumed to follow normal distribution [26]. A read is regarded as a proper read if the insert size belongs to [*μ* – 3*σ, μ* + 3*σ*] and the orientation is consistent with its mate, and is a discordant read otherwise. A read is regarded as a clipped read if it contains at least 20 unaligned bases at either end of the read, and a read is regarded as a supplementary read if different parts of the read are aligned to different regions of contigs.

The coverage-based features include standardized read coverage, fragment coverage and their deviation. The read coverage per base represents the number of reads that are mapped over that base, and the fragment coverage is the number of proper paired-end reads spanning that base. The read coverage and fragment coverage are further standardized as the ones divided by the means of the corresponding coverages of all bases across the contig or a given region.

The nucleotide variants information is extracted from BAM file with the help of samtools. metaMIC counts the number of discordant, ambiguous and correct alignments separately at each position. For each type of alignment in a contig, metaMIC will calculate the proportion of the alignment by dividing the number of this type of alignments to the total number of mapped bases across the contig, and the same for a given region.

metaMIC calculates the *k*-mer abundance difference (KAD) at each base based on the alignment of paired-end reads to contigs. The KAD value, proposed by He et al [16], measures the consistency between the abundance of a *k*-mer from short reads and the occurrence of the *k*-mer in the genome. A *k*-mer with KAD value not belonging to [-0.5, +0.5] will be regarded as an error *k*-mer, and a base is regarded as an error base if an error *k*-mer covers that base. For a given contig, metaMIC will count the number of error bases across the contig and divide it by the contig length. The proportion of error bases within a given region from a contig will be calculated in the same way.

In summary, the above these four types of features will be extracted for the whole contig (contig-based features) or a window of 100bp (window-based features). The contig-based features will be used to train a random forest to identify misassembled contigs, while the window-based features will be used as input of isolate forests to recognize the error regions containing breakpoints.

#### Identification of misassembled contigs

With the above contig-based features, metaMIC trains a random forest [35] implemented in Scikit-Learn [36] to discriminate misassembled contigs from those correctly assembled ones, where an ensemble of 1,000 trees are used. For each contig, a probability score representing the likelihood that the contig is misassembled will be output by metaMIC. The random forest model was trained on a training dataset containing contigs assembled from simulated bacterial metagenomes, whereas the ground truth misassembly labels of contigs provided MetaQUAST are used as a target for training the model. Due to the existence of strong class imbalance, we down-sampled the training dataset to obtain the same number of correct contigs paired with the misassembled contigs.

#### Localizing breakpoints in misassembled contigs

After identifying misassembled contigs, metaMIC is able to localize the misassembly breakpoints in those misassembled contigs. Firstly, metaMIC scans each contig with a sliding window of 100bp, and calculates an anomaly score for each window by employing isolation forest [37] based on window-based features to localize the error regions containing misassembly breakpoints, where the region with a higher anomaly score may be an error region; Secondly, metaMIC uses the read breakpoint ratio to recognize the exact misassembly breakpoint in an error region. Specifically, for a given predicted misassembled contig, metaMIC identifies a 100bp region with the highest anomaly score as an error region and then the position with the highest read breakpoint ratio within this window as the misassemly breakpoint. For those error regions without read breakpoints, the central position of the error region is regarded as the misassembly breakpoint.

### Evaluation of binning results

When evaluating a set of bins reconstructed from simulated microbial datasets, we use BLASTn to map each bin against the ground truth genomes used for each dataset. A representative genome of each bin is determined based on the genome which can be covered by the highest fraction of nucleotides from that bin. Then for each bin, we define the number of nucleotides in the bin that belong to the representative genome as true positives (TP). The total number of nucleotides from the bin not covered by the representative genome corresponds to the false positives (FP), and the number of nucleotides in the representative genome not covered by any contigs from that bin represents the false negatives (FN). Then the completeness, contamination and F1 score of each bin can be calculated as follows.

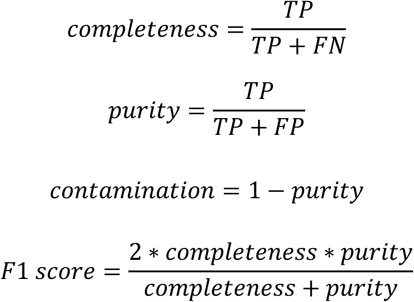

For the real metagenomics data sets where the ground truth genomes are inaccessible, we employ CheckM [38] to estimate the completeness and contamination of each bin.

## Abbreviations

MAG: metagenome-assembled genomes
bp: base pair
AUPRC: area under the precision-recall curve
KAD: *k*-mer abundance difference

